# A Bayesian Approach to Hyperbolic Multi-Dimensional Scaling

**DOI:** 10.1101/2022.10.12.511940

**Authors:** Anoop Praturu, Tatyana Sharpee

## Abstract

Recent studies have increasingly demonstrated that hyperbolic geometry confers many advantages for analyzing hierarchical structure in complex systems. However, available embedding methods do not give a precise metric for determining the dimensionality of the data, and do not vary curvature. These parameters are important for obtaining accurate, low dimensional, continuous descriptions of the data. To address this we develop a Bayesian formulation of Multi-Dimensional Scaling for embedding data in hyperbolic spaces that can fit for the optimal values of geometric parameters such as curvature and dimension. We propose a novel model of embedding uncertainty within this Bayesian framework which improves both performance and interpretability of the model. Because the method allows for variable curvature, it can also correctly embed Euclidean data using zero curvature, thus subsuming traditional Euclidean MDS models. We demonstrate that only a small amount of data is needed to constrain the geometry in our model and that the model is robust against false minima when scaling to large datasets. We apply our model to real world datasets and uncover new insights into their hierarchical structure derived from our geometric embeddings.

## 1 Introduction

Hyperbolic geometry has gained traction recently as a powerful new framework for understanding complex hierarchies in both machine learning and the basic sciences. Hyperbolic space can informally be thought of as the continuous analog of a tree, and so the exponential expansion of hyperbolic spaces allows them to capture hierarchical structure with only a few degrees of freedom. This has spurred a variety of techniques for embedding taxonomies, networks, and continuous datasets in these spaces [14, 1, 5]. For example [22] used hyperbolic embeddings to show that volatile metabolites from plants and animals conform to a low-dimensional hyperbolic geometry. It has also been shown that real world networks such as the internet possess a latent hyperbolic geometry that allows for efficient communication [1], and [10] has proposed a general framework for understanding how scale-free network topologies arise from networks being embedded in hyperbolic spaces.

Hierarchical structures are typically understood in the form of graphs, so previous representation learning studies in hyperbolic space have focused on embedding explicit networks or taxonomies [14, 15, 5, 3], where links between nodes determine their geometric similarity. However in many cases explicit hierarchical relationships are not known beforehand, and often the hierarchy cannot be decomposed cleanly into a tree-like graph [20]. Data typically instead have continuous relationships, more akin to a distance or similarity, than a binary connection. Even in studies which have worked with data in this form [9, 4], there is no clear prescription for determining the curvature or dimension of the underlying space. However, both are important geometric parameters for interpreting continuous maps obtained from discrete data. For example, dimensionality can be used to derive a minimal set of independent parameters to describe variations in the data, and curvature can act as a continuous indicator of how hierarchical the data are. This emphasizes the need for an embedding framework which can explicitly fit for the proper curvature and dimension of the hyperbolic space. In particular, complex systems typically have many degrees of freedom, but display a large scale coherence and organization which suggests the dynamics have a reduced “effective” dimension. We seek a systematic treatment of complex systems that allows us to infer their low dimensional structure, and hierarchical connections.

In this study we formulate the hyperbolic embedding problem within a Bayesian framework for Multi-Dimensional Scaling. Previous studies have investigated Bayesian MDS [16], but restricted themselves to Euclidean space. While embedding problems are typically stated as the task of minimizing some stress function, we instead formulate the equivalent maximum likelihood problem and re-interpret the stress of classical MDS [11] as a probability distribution. This allows us to incorporate hyperparameters, such as curvature, directly into the model by introducing their prior distributions. We also leverage the “Occam’s Razor” property of Bayesian statistics [12], and give a simple criteria for unambiguously determining dimension based on the evidence integral.

## 2 Hyperbolic Geometry

### 2.1 Distances and scaling

There exists much literature on hyperbolic geometry and its relevance to the study of complex networks [10]. We refer the reader to these papers for more detail, and instead focus here on the essential ideas for the present work. In the “native” coordinate representation of hyperbolic spaces the radial coordinate *r* of a point is equal to its distance from the origin. In this representation we can compute the distance *x* between 2 points with angular separation Δ*θ* by the hyperbolic law of cosines

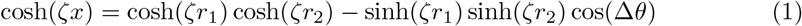

where *ζ* defines the sectional curvature *K* of the space as *K* = −*ζ*^2^. Note that as *ζ* → 0 this reduces to the Euclidean law of cosines, as expected.

Although the straightforward scale invariance of flat spaces is lost, Hyperbolic spaces still possess a more subtle form of scale invariance which our MDS algorithm will exploit. Consider a joint rescaling of the coordinates *r* → *λr* and curvature *K* → *λ*^-2^*K*. By Eq. 1 the distance must then be rescaled as *x* → *λx*. Thus, unlike Euclidean spaces which can be rescaled simply by scaling the coordinates, scaling hyperbolic spaces must also be accompanied by a rescaling of the curvature. Furthermore, we see that up to an overall scaling of distances, a hyperbolic space with unit curvature and maximum radius *R_max_* is equivalent to a space with unit radius and curvature 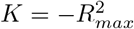. This allows us to modulate the curvature of our space at fixed radius simply by rescaling our distance matrix, a fact we will exploit in section 3 in order to adaptively fit for the curvature of our embedding space.

### 2.2 Embedding coordinates

There are many equivalent coordinate systems to describe hyperbolic spaces [10]. Although “compact” projections such as the Poincaré or Beltrami-Klein representation can be intuitive for visualization, we follow [15] who found that the Lorentz model is significantly better behaved computationally. In this model a *D* dimensional hyperbolic space is represented by a future facing space-like sheet in a *D* +1 dimensional Minkowski space-time. I.e. the set of points satisfying

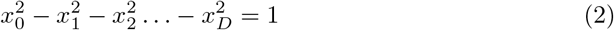

If we introduce a metric tensor *g_αβ_* = Diag(1, −1, −1,…, −1) on the D + 1 dimensional space, we can compactly define our hyperbolic subspace by the constraint equation

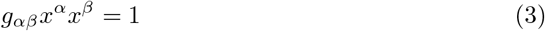

where greek indices *α* and *β* run from 0,…, *D* and we have employed the Einstein summation convention that repeated indices are summed over. We can compute the hyperbolic distance between any 2 points satisfying these constraints as

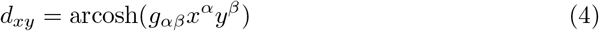

Computationally, we take the *D* space-like components 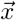 of the coordinates as our free parameters, and compute the time-like component according to the constraint 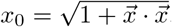.

Note that these coordinates are for a space with unit curvature *K* = −1. As we will see in the next section, our embedding model will fit for the maximum radius of the distribution of points in this space. Once the embedding is complete, we can rescale distances and coordinates and reinterpret the model as having a different curvature, but for computational simplicity it is preferred to perform the embedding itself at fixed curvature.

## 3 A Bayesian model for MDS

We now turn to describing our Bayesian model for hyperbolic MDS, and inferring dimension. We demonstrate that our model is both efficient and scalable. Only a small number of points is required to correctly infer the curvature of the space, and our modelling of embedding uncertainty makes the optimization robust against false minima. Finally, we present an iterative algorithm for effectively scaling the optimizer up to large datasets.

### 3.1 The likelihood function

Given a matrix *δ_ij_* of distances (dissimilarities) between data points, Multi-Dimensional Scaling seeks an embedding of points 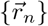 in a geometric space whose distance matrix *d_ij_* matches the dissimilarity matrix as closely as possible [11]. This is formulated by defining a *stress* function which is minimized when the matrices are exactly equal.

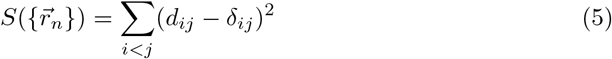

As has been pointed out in previous Bayesian studies [16], minimizing Eq. 5 is equivalent to finding the maxmimum likelihood of an associated Gaussian likelihood function. Taking this perspective, we seek to construct a generative stochastic model for the data such that the posterior distribution 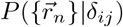 is maximized exactly when the stress is minimized.

By analogy with generative models for linear regression, we model our data dissimilarites as being generated directly from an underlying geometric model by some stochastic process that introduces white noise to the system. Thus we write.

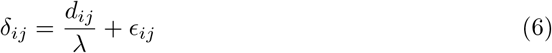

Where the 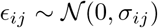 are independent and normally distributed random variables, but with possibly differing variances, and *λ* is a global scale parameter. Without loss of generality we normalize ***δ*** to have a maximum value of 2 (i.e. unit radius), so that we can view *λ* as setting the maximum radius of the embedding with unit curvature *K* = −1. Equivalently, as discussed in Sec. 2 we can interpret λ as setting the curvature of the interior of the unit sphere to be −*λ*^2^. From the form of the distribution for *ϵ_ij_* we can write the conditional distribution 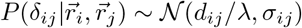. Taking the product over all pairs of points gives the likelihood of the parameters given the dissimilarity matrix

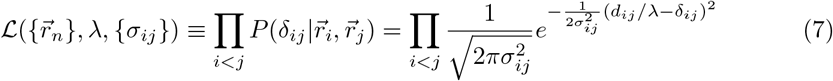

For given values of the parameters λ and constant {*σ_ij_*} the maximum likelihood solution of Eq. 7 is equivalent to the minimum of Eq. 5. However it is unclear what the optimal values of these parameters should be, and in the case of *λ*, the actual value of the parameter has geometric implications for the interpretation of the model. Therefore we take a Bayesian approach and fit for all parameters simultaneously by introducing priors over 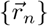, *λ*, and {*σ_ij_*} to compute the posterior

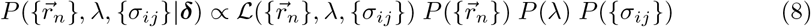

### 3.2 The posterior distribution

We start by choosing prior distributions for our parameters which appropriately regularize them without being too restrictive. For our embedding coordinates we would like something like a harmonic oscillator potential that posesses spherical symmetry and prevents points from escaping to extremely large radii. Though we could implement this directly with a normal prior on the radial coordinates, this is a complicated function of the lorentzian coordinates and could impair the speed and stability of the code. Instead, we use the fact that *λ* controls the size of the space, so putting a normal prior on *λ* will have the same effect and is significantly simpler. Note that the log-likelihood scales with the number of points as ~ *N*(*N* – 1)/2, while *λ* ~ *N*^0^. Thus as the number of points in our embedding increases the prior on lambda becomes negligible relative to the likelihood. To remedy this we multiply the log-prior on *λ* by *N*(*N* – 1)/2 so that it scales with the likelihood. With this, we can simply leave a flat prior on the Lorentzian coordinates. Although there is a Jacobian distortion of the flat prior when transforming back to the native hyperbolic space, the effect is negligible and the embedding results are unaffected by it.

For the embedding variances we reduce the number of parameters by introducing an uncertainty *σ_n_* associated to each data point 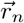. We then compute the uncertainty of the distance between points *i* and *j* as

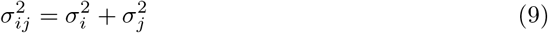

From a physical perspective, minimizing the stress is equivalent to finding the lowest energy configuration of a collection of points fully connected by springs of equilibrium lengths given by *δ_ij_* and stiffnesses 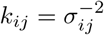. Thus the interpretation of Eq. 9 is that each point has a characteristic stiffness 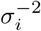, and each pair of points is connected by their 2 springs in series. Physically, this model allows subsets of points which are well fit relative to each other to condense into high-stiffness/low-uncertainty clusters, while poorly fit points have low stiffness and can still easily explore the space. Not only does this help the optimizer (Fig. 1 left panel), but it also allows us to identify if points have gotten caught in a false minima and help guide them out of it (Fig. 1 middle and right panels). This allows us to iteratively handle problems with false minima when scaling up to large datasets. We complete our model by putting an inverse-gamma prior on each *σ_i_*, as a standard semi-informative prior for gaussian variances. We found that our results are not sensitive to the choice of the prior on *σ*, so long as it is not too restrictive.

**Figure 1.**
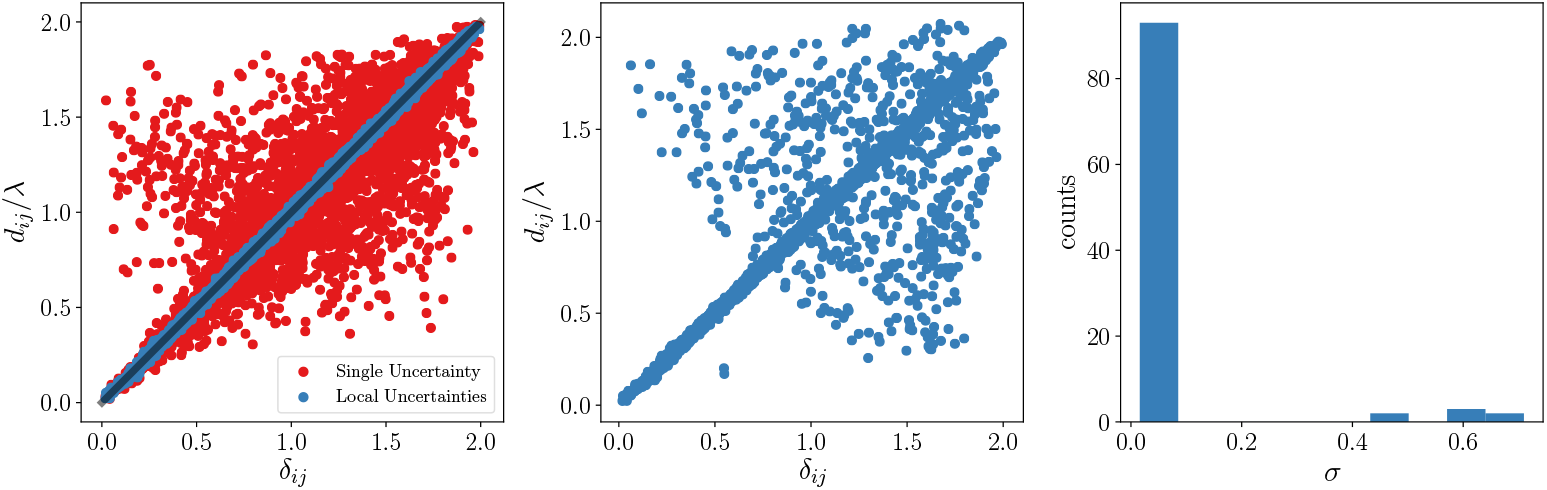
Uncertainty Modeling. Left: “Shepard Diagrams”, plot of *d_ij/λ_* versus *δ_ij_*, for an embedding in 2D with a single global uncertainty *σ* in red, and an embedding with individual uncertainties *σ_n_* for each point in blue. The optimal solution should coincide with the line of slope 1 show in grey. Middle: An embedding with local uncertainties caught in a false minimum. Most points are well fit to each other, with a few poorly fit points responsible for the observed scatter. Right: Distribution of *σ* values for the false minimum embedding. We can clearly identify a stiff and loose cluster of points. We can randomize the coordinates of points with high *σ* and re-run the optimizer until all points are at the optimal solution.

Putting all this together, we can write the posterior distribution as

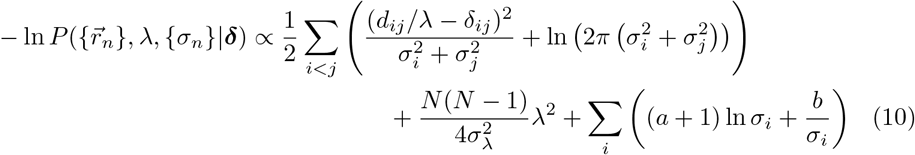

This represents the objective function which we seek to minimize by our embedding. The crucial point is that the embedding distance matrix *d_ij_* is computed with respect to a hyperbolic metric, and the embedding coordinates 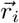 are the lorentzian coordinates discussed in section 2.2. We chose *σ_λ_* = 10 and *a* = 2, *b* = 0.5, though we have confirmed that the embedding results are insensitive to the choice of hyper-parameters. We minimize Eq. 10 using an L-BFGS algorithm implemented in the Stan statistical package [2], distributed under a BSD license. We deal with the singularity at *λ* = 0 by imposing *λ* > 0.001. We confirmed that this value is small enough that the hyperbolic law of cosines with the corresponding curvature gives indistinguishable results from the Euclidean law of cosines, thus this lower bound is small enough to cover the euclidean case.

### 3.3 Synthetic data results

We first test our method on synthetic distance matrices produced by randomly generating points uniformly in hyperbolic spaces out to some maximum radius. We add noise of magnitude 0.05*R_max_* to each distance matrix to simulate a more realistic dataset.

To test our method’s ability to fit for the underlying curvature of the space we generate points in 3 dimensional spaces with *K* = −1 out to varying maximum radii. We then rescale all of the distance matrices so that their maximum distance is 2 (i.e. radius of 1), and we are thus ignorant as to the true radius of the data. With this rescaling the value of λ predicted by the simulation is exactly the predicted *R_max_*, and we can compute the model curvature as *K_model_* = −*λ*^2^ (recall the discussion in Sec. 2.1). We see in panel A of Fig. 2 that we are able to effectively fit for the correct curvature with as few as 25 – 50 points. In orange we show the results of embedding data generated in a flat space with noise. Our model correctly predicts curvatures close to 0, and the embedding matrix matches *δ_ij_* almost perfectly. Thus our method *subsumes* traditional MDS methods which only operate in flat spaces. We also show in panel B how the variance in *λ_model_* with respect to different random seeds converges as a function of the number of points.

**Figure 2.**
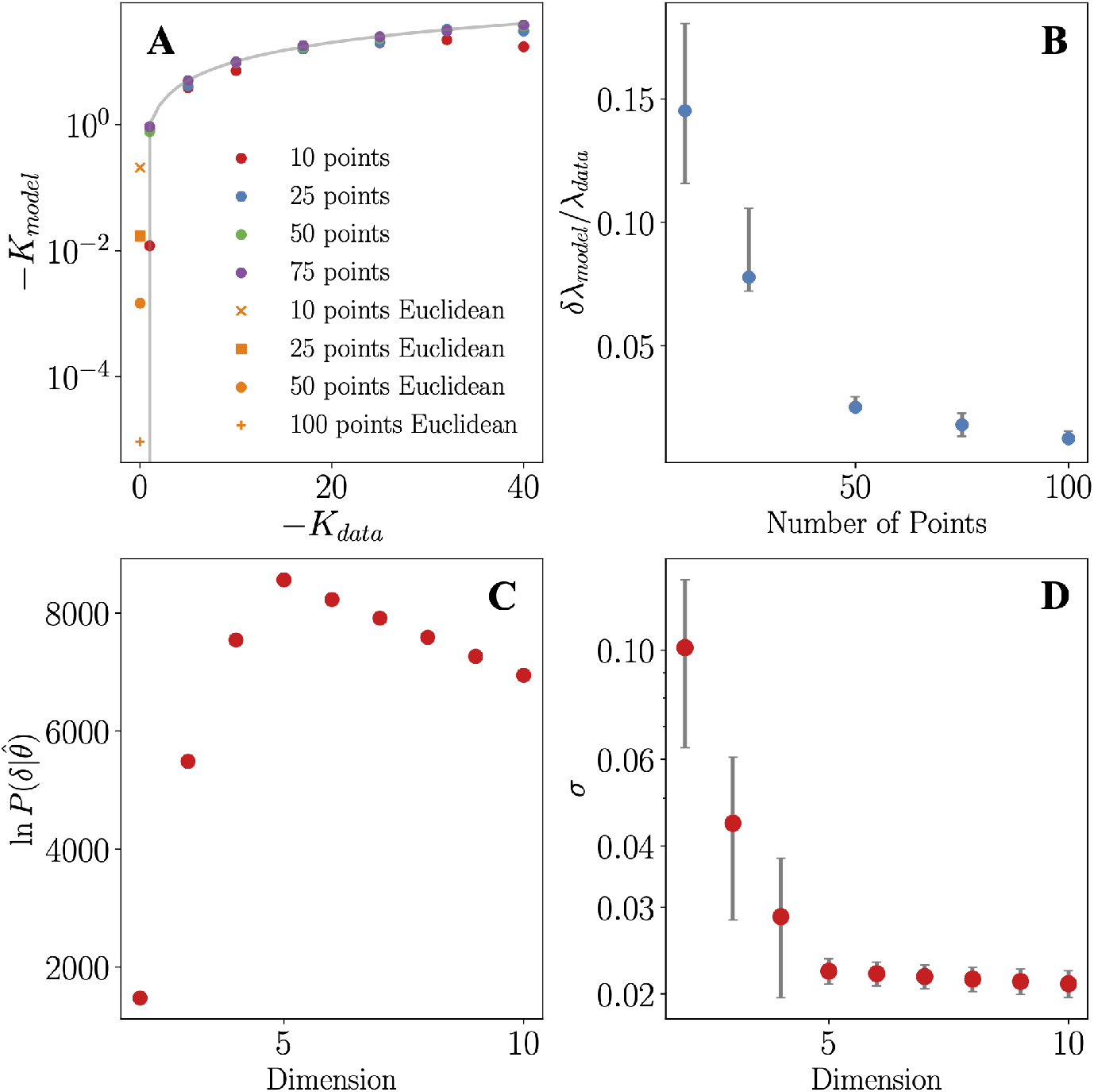
Curvature and Dimension. A: Predicted curvatures *K_model_* plotted against actual curvatures *K_data_* for synthetic data. *K_model_* = *K_data_* line is shown in grey. Predicted curvatures for Euclidean data are shown in orange. B: fractional error in *λ_model_* (*λ_data_* = 5) as a function of the number of points. Errors bars show uncertainty with respect to simulating multiple times with different random seeds. C: Bayesian Information Criteria versus embedding dimension evaluated on a synthetic dataset of true dimension *D* = 5. D: Mean *σ* for the same embeddings.

One advantage of a Bayesian approach to MDS is that we can employ Bayesian model selection techniques to determine the underlying dimension of the space. Let 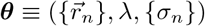 denote the set of all parameters. Naively, one could compute the likelihood 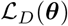 of an embedding in dimension *D* and choose the dimension which maximizes this value [11], however this does not account for volume effects introduced by having a different number of parameters in embeddings of different dimension. Instead one would like to compare models by computing the evidence [12], which integrates out all parameters and defines the Bayesian Information Criteria (BIC) [19] as:

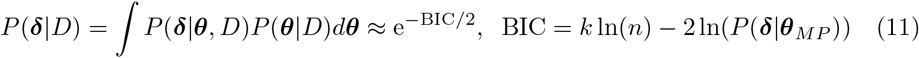

Where *k* is the number of parameters in the model, *n* is the number of observations, and ***θ**_MP_* are the values of the parameters that maximize the posterior. This approximate expression for the evidence integral is obtained by evaluating the integral according to the saddle point method. Since the posterior is invariant to rigid rotations of the configuration of points there are *D*(*D* – 1)/2 “redundant” transformations which do not change the probability distribution. We exclude these rotational degrees of freedom when computing the number of model parameters so *k* = *N*(*D* + 1) + 1 – *D*(*D* – 1)/2.

To test our method’s ability to infer the underlying dimension of the space we generate 100 points in a 5 dimensional hyperbolic space, and embed the resultant normalized distance matrix across a range of dimensions. We compute the evidence for all embeddings and plot the result in panel C of Fig. 2. For *D* < 5 the evidence decreases due to the poor quality of fit, while for *D* > 5 the evidence decreases since the quality of fit remains constant but the number of parameters increases. Penalizing additional parameters which do not aid the model is one of the principle features of using the evidence for model selection. Thus we are able to correctly identify *D* = 5 from the maximum of the evidence.

Alternatively, we can analyze the distributions of the *σ* values of the embeddings, shown in panel *D* of Fig. 2. There is a clear “elbow” at *D* = 5 where the mean and standard deviation of the distribution drops dramatically. Adding more dimensions gives little to no improvement so we can still identify the correct dimension as *D* = 5. When working with real datasets, however, this elbow can often be less clear so we use the BIC going forward.

### 3.4 Techniques for Embedding Large Datasets

The highly non-linear nature of hyperbolic spaces which endows them with the geometric power to capture complex hierarchical relationships, also makes them very difficult to deal with numerically. We describe here an iterative procedure for scaling up the BHMDS algorithm to embed arbitrarily large datasets.

The primary difficulty with large datasets is that for random initial conditions the gradient of each contribution to the cost function tends to be very large. Summed over ~ *N*^2^ pairs of points this can easily lead to an overflow. Our solution is to allow the simulation to find the “optimal” initial condition so that initial gradients are minimal. To do this, suppose that a subset *N_seed_* < *N* points have been embedded in a *D* dimensional hyperbolic space and we would like to add 1 more point to the embedding. Since *N* > *N_seed_* > *D* the distances between the new point and the existing *N_seed_* points are in theory enough to constrain the position of the new point. So, we freeze the *N_seed_* points in place and find the position of only the new point based on its distances to the *N_seed_* frozen points. Since the data distance matrices are noisy we expect this to only be approximate, so we think of this procedure as finding the optimal initial condition for the new point. We finish by unfreezing and fully coupling all points as if it were standard MDS and continue optimizing the cost function until convergence.

The essence of our iterative algorithm is to scale this procedure up by adding points in batches of 100 at a time, instead of one at a time. We do this as follows:

- Start with random subsample of *N_seed_* points out of the full N points and embed them according to the standard BHMDS algorithm. We typically take *N_seed_* = 300, but any number small enough that it can be manageably embedded will work.
- Select 100 new points which have not been embedded yet. With the *N_seed_* points frozen in place we “initialize” each of the 100 new points individually in the manner described above.
- Unfreeze all points and optimize the full cost function of *N_seed_* + 100 points until converged.
- Update *N_seed_* = *N_seed_* + 100 and continue adding points in batches of 100 until *N_seed_* = *N*

The procedure is flexible in the explicit choices of 300 points for the initial embedding and adding points in batches of 100. We found success embedding up to 10, 000 points with this method, though we have not yet fully probed the upper limit of what this algorithm can embed.

## 4 Illustrative Examples

We now consider two example problems with real world data from broadly different fields to illustrate the power and versatility of the Bayesian hyperbolic MDS. The first example deals with how transportation networks organize themselves into hierarchies in complex systems, and the second investigates the hierarchical nature of viral evolution from a geometric perspective.

### 4.1 Transportation networks as hyperbolic systems

As our first example we ask how transportation networks in complex systems may be analyzed using the tools of hyperbolic geometry. Networks such as vasculature in the body for transporting blood, or roadway networks for transporting cars, display a complex branching structure indicative of some sort of hierarchy: large fast moving vessels branch off into smaller slower moving ones [8, 7, 21]. How might we concretely map out this hierarchy using the tools of hyperbolic geometry?

To do this we take the roadway network of San Diego as our example system. Although San Diego is a sprawling metropolitan area, there is local wisdom that it takes roughly 20 minutes to drive between any 2 points in the city. One can confirm on Google Maps that the travel time between 2 points scales only weakly with their physical distance. This is similar to the property of hyperbolic spaces that at large radius and curvature, the distance between 2 points depends only logarithmically on their angular separation. Taylor expanding Eq. 1 for large *ζr* gives

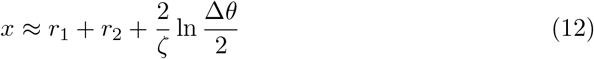

At large curvature most of the volume in a hyperbolic space is at large radii, so for a uniform distribution of points out to a large maximum radius we expect most points to have *r* ~ *R_max_*, and thus expect distances between pairs of points to be nearly constant. To characterize the transportation network of San Diego we propose to make a map of the city where distances between points are the time it would take to travel by car between them; we refer to the embedding informally as the “time space” representation of the transportation network. Based on the above analogy we theorize that this map will be characterized by a hyperbolic geometry which will allow us to map out the hierarchies present in the network. To do this we curated a dataset of 100 points sampled throughout the city of San Diego (see Fig. 3 middle panel). 40 points were used to sample the highway network as evenly as possible, and the remaining 60 were used to randomly sample addresses from the rest of the city. We used Google Maps [13] to estimate the driving time between every pair of points. It is then straightforward to find an embedding of this distance matrix using both the Bayesian hyperbolic-MDS method presented above as well as traditional Euclidean MDS for comparison. From a Shepard diagram analysis alone (Fig. 3 left panel) we see that the Hyperbolic embedding gives a much better fit than Euclidean. This is further confirmed by the fact that the BHMDS algorithms predicts a maximum radius of *λ* = 4.03. If the data were in fact Euclidean, the algorithm would have predicted *λ* ≪ 1 (see Fig. 2).

**Figure 3.**
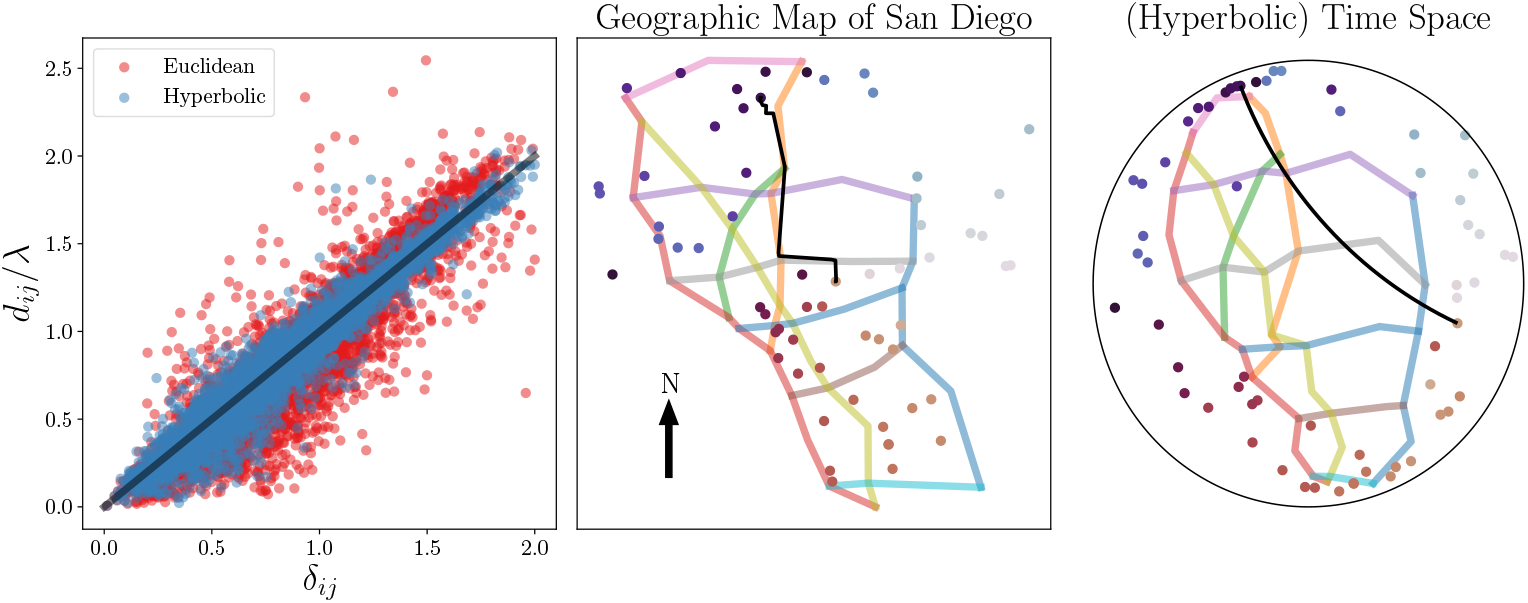
Transportation Networks. Left: Shepard diagram for both traditional Euclidean (red) and BHMDS (blue) embeddings of the travel times distance matrix. Middle: physical locations of city points on a geographic map of San Diego. Points from the highway network are connected, and local addresses are colored by their polar angle from the center of the city. Google Maps driving directions are shown between a pair of randomly selected points. Right: Hyperbolic embedding of the travel time matrix in the Poincare plane. Colors are preserved from the middle panel for comparison between the two maps. The hyperbolic geodesic is plotted between the 2 points which we show the driving directions between in the middle panel.

In the right and middle panels of Fig. 3 we plot the hyperbolic embedding of the points based on their pairwise travel time distances next to their positions in physical space, with the highway network highlighted and connected. We can immediately see the hierarchical structure of the network mapped out: highways get pushed to the origin, representing the root node of the hierarchy, while surface street addresses get pushed to large radii and act as leaf nodes. Note that the angular coordinate of the points are roughly preserved by the embedding. The interpretation is that to get from point A to point B in the city one first travels from point A to the highway, and then exits from the highway to reach point B. This is easily confirmed by studying the Google Maps driving directions between points in the city: unless 2 points are extremely close to each other the directions will route you to the nearest freeway to reach your destination, which almost never coincides with the physically shortest path between the points. This observed hierarchy also explains our initial observation that it takes ~ 20 minutes to drive between any two points in San Diego. The majority of driving time is spent getting to and from the highway, whereas the majority of the distance traveled is covered in a short amount of time on the freeway, which explains the weak scaling of travel time with distance [8]. This is similar to a hyperbolic space in which the majority of the distance contributed to the geodesic between 2 points is from traversing to and from the origin, while the most angular displacement happens very rapidly with a quick turn at small radii.

Finally we observe how geodesics in the hyperbolic space relate to shortest time driving directions in the geographic map of the city. In the middle panel the driving directions between 2 randomly chosen points take you west along the grey highway and then north along the orange (note that this path is markedly different from the Euclidean geodesic on the geographic map). The geodesic between the corresponding points in the hyperbolic space run adjacent to the hyperbolic representations of the same highways. This presents an intruiging geometric approach to search for the appropriate highways (i.e. root nodes) to route you from one point to another in a city.

Future work could extent our approach of embedding a matrix of travel times to study other transportation networks such as vasculature or protein transport in cells. One could also study how traffic jams and other dynamical effects affect the geometric representation of the space of travel times. It is possible that the phase transition like “freezing” of traffic in a jam [6] can be associated with a breakdown of hierarchy in the network, since in a jammed state highways no longer act as faster root nodes, and could potentially be interpreted as a transition between hyperbolic and Euclidean geometries.

### 4.2 The geometry of evolution

For our second example we analyze the hierarchical nature of evolution through the lense of hyperbolic geometry. The conception of evolution as a vast branching tree was immortalized early on by Darwin with his depiction of a “tree of life” [20]. Based on this analogy we theorize that hyperbolic geometry can be used to effectively map out the evolution of new species. To study this quantitatively we use the database of COVID-19 gene sequences provided by the NCBI [18]. We seek to map out the genetic evolution of COVID-19, and provide a geometric basis for the identification of new strains.

To compile our dataset we take a random sample of 1000 COVID sequences sampled uniformly in time between Jan 1st 2020 and Oct 1st 2021. We measure distance between sequences by counting the number of nucleotide positions in which two gene sequences *disagree*, also known as the Hamming distance. We embed this distance matrix over a range of dimensions using the large scale embedding algorithm presented in section 3. The resultant embeddings are strongly hyperbolic, with a predicted maximum radius of *λ* = 4.97 in a 3 dimensional hyperbolic space. We can also confirm this by comparing the shepard diagrams of hyperbolic and Euclidean embeddings in the optimally predicted dimension. From the inset of the middle panel we can clearly see the hyperbolic embedding gives a better fit to the data. For plotting purposes we project out one of the dimensions of the 3D embedding and plot the results in the Poincare plane in the left panel of Fig. 4. A temporal hierarchy is immediately revealed by the embedding: in the right panel we see that the hyperbolic embedding radius scales with the date that the sequences were collected. This suggests that we are seeing an evolutionary hierarchy unfolding in time in hyperbolic space.

**Figure 4.**
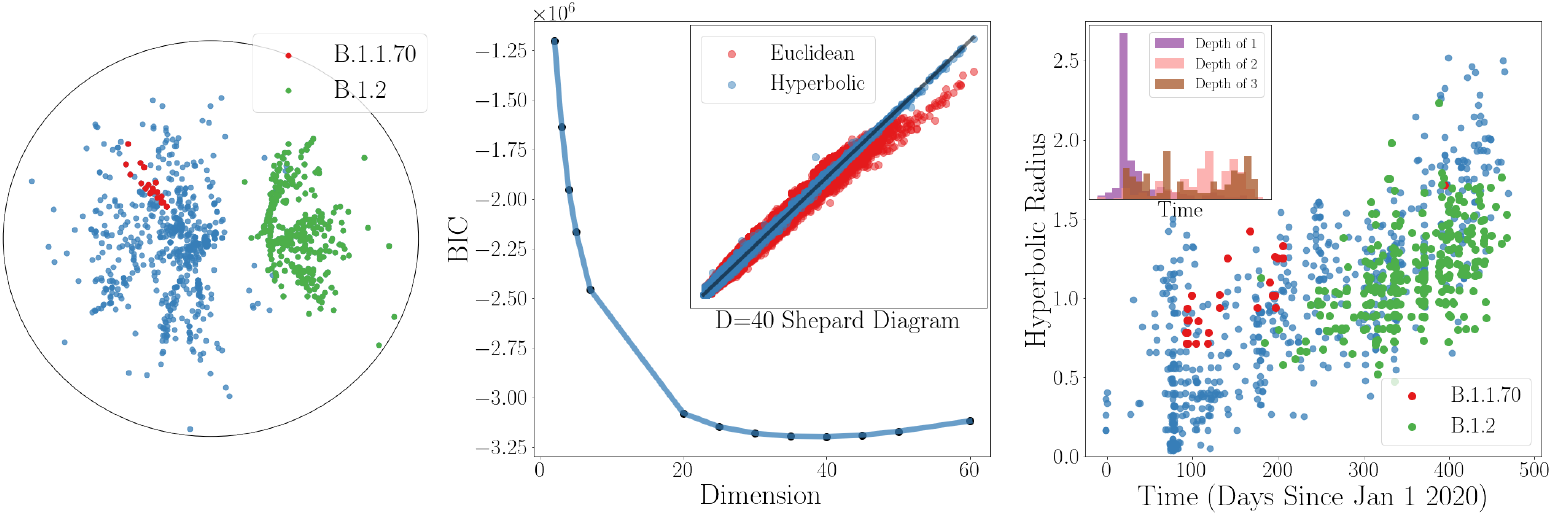
Viral Evolution. Left: Embedding of covid sequences in 3D hyperbolic space with one of the dimensions projected out. Example PANGO lineages are shown in red and green. Middle: BIC curve for embeddings across a range of dimensions. Middle Inset: Shepard diagrams for hyperbolic and Euclidean embeddings at the optimal dimension. Right: Hyperbolic radius of covid sequences plotted against their date of collection. Right Inset: Histogram of collection dates of covid sequences of different hierarchical depth as predicted by the Pangolin algorithm

We can compare our results to the Phylogenetic Assignment of Named Global Outbreak Lineages (PANGOLin) [17] labels for the sequences. PANGOLin provides a system for labeling new lineages of covid strains as they emerge, and also gives a rough measure of the hierarchical “depth” of each sequence (i.e. how many lineages down from the “root” sequence). In the left panel we can see that different Pango lineages trace radial arcs in the hyperbolic space, suggesting that the branching structures in the embedding correspond to the evolution of different lineages. In the inset histograms in the right panel we see that the PANGOlin predicted hierarchical depth of each sequence is only a weak predictor of the collection dates. This suggests that the our continous geometric model is a more robust indicator of the hierarchical structure than the discrete PANGOlin predictions of depth.

Finally we can analyze the BIC to determine the minimal number of degrees of freedom to characterize variations in covid gene sequences. In the middle panel of Fig. 4 we see the BIC is minimized around a dimension of *D* ~ 40, which we identify as the minimal dimension. While this may seem like a large dimension, we must keep in mind that each sequence has ~ 50,000 nucleotides in it, so this represents a vast reduction in the number of degrees of freedom. Potential future work would identify each of these reduced degrees of freedom with coarse grained biological functions.

## 5 Discussion, ethics, and future work

We have presented a Bayesian method for embedding data in hyperbolic spaces, with a novel approach to uncertainty modelling, as well as improved techniques for inferring the curvature and dimension of the underlying space. We established through tests on synthetic datasets that our method is both accurate and efficient: the algorithm consistently reconstructs the data in space with high fidelity, and can correctly infer the geometric parameters of the space with very little data. We emphasize a novel feature of our model is its ability to both fit data to geometry, through MDS embedding, and the ability to fit geometry to data, through the Bayesian inference of geometric hyper-parameters. On real datasets from complex systems, our hyperbolic methods show vast improvements over Euclidean embeddings and uncover novel insights about the hierarchical nature of the data. We hope our analyses of these example problems will spur more in depth investigation of the geometry of these systems. We also emphasize that embeddings allow us to infer the underlying hierarchy in the data in a continuous manner, and thus can afford more power and flexibility than discrete hierarchical clustering algorithms.

Our model is limited in its assumption of spaces with uniform curvature and integer dimension. Modeling spaces with uniform curvature amounts to the assumption that the underlying hierarchy can be cleanly decomposed into a b-ary tree with constant branching factor. This is, however, an over simplification of real world hierarchies. One could address this by exploring generalizations of the model to spaces with non-uniform curvature, such as curvature varying with radius, or different curvatures along different axes. By modeling our data with a space of uniform dimension we ignore the possibility of having different length scales characterized by a different number of degrees of freedom. For example the local structure of the data could be characterized by high dimensional noise, while the global correlations could constrain the data to a lower dimension at large lengthscales. One could address this in future work by incorporating “compact hidden dimensions” akin to in string theory.

The concept of hierarchy has long been used as a justification for the unequal treatment of people on the basis of race, gender, and wealth. A possible danger of a study such as this is that malicious practitioners may apply such hierarchical models in an anthropological setting to promote policies or theories of society that are fundamentally anti-humanitarian. We urge practitioners to exercise strong caution if these methods are used to look for structure in datasets pertaining to academic or economic standing, race, gender, or urbanization. In particular, practitioners must investigate the socio-economic factors relevant to the system at hand if one is to use these methods on such datasets. Otherwise, one runs a strong risk of ascribing erroneous and harmful interpretations to the data.

